# A New Uranium Bioremediation Approach using Radio-tolerant *Deinocoocus radiodurans* Biofilm

**DOI:** 10.1101/503896

**Authors:** T. Manobala, Sudhir K. Shukla, T. Subba Rao, M. Dharmendira Kumar

## Abstract

*Deinococcus radiodurans*, is the most radiation tolerant organism ever known, it has gained importance in recent years as a potential candidate for bioremediation of heavy metals, especially radioactive ones. This study investigates the efficiency of a recombinant *D. radiodurans* (DR1-bf^+^) strain with an ability to form biofilm for uranium remediation. The modified Arsenazo III dye method was used to estimate the uranium concentration. Uranyl nitrate aqueous solution is generated during the operation of nuclear fuel reprocessing. The *D. radiodurans* biofilm (DR1-bf^+^) grown in the presence of 20 mM Ca^2+^ showed remarkable ability of uranyl ion removal. DR1-bf^+^ (+Ca^2+^) biofilm removed ∼75 ± 2% of 1000 mg/L uranium within 30 minutes post treatment from uranyl nitrate aqueous solution. U removal rate was also found to be directly proportional to biofilm age. This study discusses the ability of *D. radiodurans* biofilm in uranium removal.

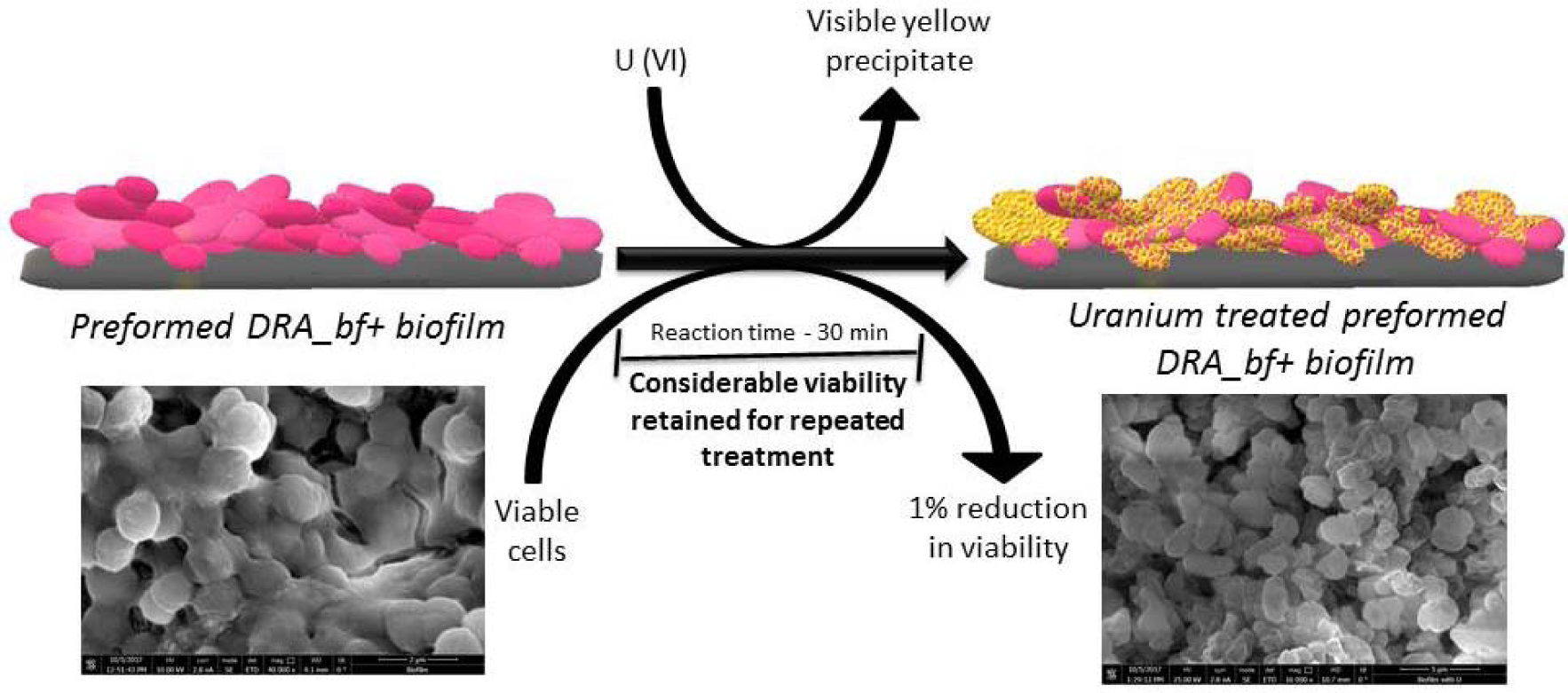
Graphical abstract.

## 1. Introduction

In a natural environment, almost 99% of all microorganisms live as surface-attached communities known as biofilms (Costerton et al. (1987). A biofilm is an assemblage of microbial communities, in which the microbial cells are irreversibly attached to a surface and embedded in an extracellular polymeric matrix (EPS) produced by these cells (Donlan, 2002; Das et al., 2012). Biofilms apart from enhancing the organism survival, improve the rate and extent of contaminant transformation as compared to that of pure and planktonic cultures (Mangwani et al., 2014). Remediation using microorganism (termed as bioremediation) is an emerging *in situ* technology for the clean-up of environmental pollutants. The economical factor and inefficiency of some physico-chemical remediation methods has made this microbiological treatment method, an improved and preferred alternative (Paul et al., 2005).

Heavy metals are released into the environment through various processes, leading to contamination of air, water, and atmosphere (Akpor and Muchie, 2010). These heavy metals need to be converted from one form to less hazardous form or its bioavailability should be decreased (Wall and Krumholz, 2006). Among the many heavy metals, naturally occurring and comparatively less abundant uranium has drawn attention due to the surge in anthropogenic activities involving uranium and its potential hazards due to an accidental release into the environment. Nuclear fuel reprocessing generates radioactive uranyl nitrate [UO_2_(NO_3_)_2_·6H_2_O] (Baumgärtner, 2009). In aqueous medium, uranium exists as uranyl ion (UO_2_^2+^) i.e. UO_2_^+2^ is highly soluble and mobile in the subsurface environments and could be hazardous being a heavy metal and radioactive (Shukla and Rao, 2015). The intake of these ions can cause detrimental health hazards (Keith et al., 2013). The chemical methods used for uranium removal have their own limitations due to very high cost and low feasibility for *in situ* remediation (Kulkarni et al., 2013).

Bioremediation of uranium using microbes can be a very promising and more efficient approach as microbes are nature’s creative recyclers (N’Guessan et al., 2008). Biofilms are ideal for bioremediation purpose as they need not to be separated from the bulk liquid waste, thus making the whole process more economical and feasible (Shukla et al., 2017). Biofilm mediated bioremediation of UO_2_^2+^ would be an ideal process by a radio-tolerant microbial biofilm to ease its further downstream processing. Also conversion of aqueous U to insoluble uranyl phosphate precipitates are difficult for reoxidation in contrast to the ‘reduced’ uranium mineral like uraninite which has the tendency to reoxidize back to more soluble U (Kulkarni et al., 2013). In such cases, inherently radio-tolerant bacterium *Deinococcus radiodurans* can be of use. *Deinococcus radiodurans* has an extraordinary capability of surviving under high radiation stress (Battista, 1997), even under low nutrient conditions (Shukla et al., 2014). The bioremediation potential of *D. radiodurans* has gained importance in recent times. Engineered *D. radiodurans* cells have been used to detoxify mercury, degrade toluene (Brim et al., 2000) and chromium reduction (Brim et al., 2006). However, use of planktonic cells for bioremediation of heavy metals makes the downstream process costly or less efficient.

A genetically modified strain of *D. radiodurans,* harboured with a constitutively expressing *gfp* plasmid (GenBank accession no. KF975402), was used in this study and found to be a biofilm forming strain (Shukla and Rao, 2017a). We have investigated whether this biofilm forming strain of *D. radiodurans* (hereafter denoted as DR-bf^+^) can be used for the biofilm-mediated uranium bioremediation from the uranyl nitrate aqueous solution.

## 2. Materials and Methods

### 2.1 Microorganism and culture conditions

*Deinococcus radiodurans* R1 wild type (DR1-WT) and *gfp-*harbouring, kanamycin resistant, biofilm forming strain of *D. radiodurans* (DR1-bf^+^) was used in this study. TGY medium consisting of tryptone (5 g/L), yeast extract (3 g/L), glucose (1 g/L) was used as growth medium. The DR1-bf^+^ bacterium was grown for 48 h in TGY medium at 30°C in an orbital shaker at 100 rpm until mid-log phase. Kanamycin antibiotic was added in the final concentration of 5µg/mL for the growth of *D. radiodurans* (DR1-bf^+^). For most of the experiments mid-log phase cultures were used.

### 2.2. Biofilm estimation using crystal violet assay

Biofilm assay was performed in 96 well microtiter plates as per the modified classical crystal violet assay (Shukla and Rao, 2017b). Late log phase *D. radiodurans* culture (grown for 48 h) were diluted in sterile TGY medium in 1:40 ratio and 200 µl of this diluted culture was transferred to the pre-sterilized 96 well flat bottom polystyrene microtiter plates. After the prescribed time as per the experimental plan, planktonic cells were aspirated and the biofilm grown in 96 well plate was washed twice with 200 µl of sterile phosphate buffer saline (PBS), air dried and stained with 0.2% crystal violet solution. For dissolution of bound crystal violet, 95% ethanol was used. Biofilm growth was estimated in terms of absorbance at 570 nm using a multimode micro-plate reader (BioTek, USA).

### 2.3 Fluorescence microscopy study

The biofilm samples were grown in petri dishes for fluorescence microscopy study. The cells were grown in TGY media at 30°C and 150 rpm in 15 mL falcon tubes. 100 µl mid log phase planktonic cells of DR1-bf^+^ (with OD of 0.7) were transferred to petri dishes (35 mm) with 4 ml of TGY media in the presence of calcium (20 mM) and required amount of kanamycin antibiotic with a final concentration of 5 μg/mL. The used up medium was aspirated and the biofilm was rinsed with sterile PBS to remove loosely attached cells. The petri plate was viewed under a fluorescence microscope (Carl Zeiss Axioscope I, Germany) equipped with a fluorescence illumination system and with appropriate filter sets. Images were obtained from randomly selected fields using 20X objective.

### 2.4 Biofilm growth of DR1-bf^+^ for uranyl nitrate treatment

The biofilm forming strain of *D. radiodurans* (DR1-bf^+^) was used in this biofilm-mediated bioremediation study (Shukla and Rao, 2015). The cells were grown in TGY media at 30°C and 150 rpm in 15 mL falcon tubes. 100 µl mid log phase planktonic cells of DR1-bf^+^ (with OD of 0.7) were transferred to petri dishes (35 mm) with 4 ml of TGY media (with or without 20 mM Ca^2+^) and kanamycin with a final concentration of 5 μg/mL. These plates were incubated in the static condition at 30°C for the required period of time and the used up media was replaced after every 48 h. After given time period, the planktonic cells were aspirated and biofilm was gently washed twice with PBS, before the treatment with filter sterilized uranyl nitrate solution (pH 3.61).

### 2.5 Uranium estimation protocol Arsenazo III dye method and by Inductively Coupled Plasma-Atomic Emission Spectroscopy (ICP-AES)

Uranium in the aqueous sample was estimated by modified protocol of Arsenazo III method. In this study, the standard arsenazo dye method (Savvin, 1961) was modified for the uranium estimation. Briefly this method involves simple treatment of the uranium solution with a buffer Potassium Hydrogen Pthalate buffer (1.5 mM PHP in 0.025 N HCl) and 0.1% Arsenazo III dye. To 1 ml of PHP buffer, 900 µl of the sample and 100 µl of Arsenazo III reagent was added. It was mixed by vortexing and absorbance was measured at 653 nm (absorption maxima for the U-Arsenazo III complex) after 5 minutes. The change in the color is directly proportional to the amount of uranium present in the given solution. Suitable dilution factors and arsenazo dye concentration were used to estimate uranium in wide range. The samples were also analyzed for uranium estimation using Inductively Coupled Plasma-Atomic Emission Spectroscopy (ICP-AES, model: Ultima-2, Horiba JovinYvon, France) to substantiate the modified Arsenazo III dye method. Two ml of uranyl nitrate sample was collected and taken for ICP-AES analysis. Standard graph was plotted for both modified Arsenazo III dye method and ICP-AES to validate the Arsenazo III dye method (Savvin, 1961).

### 2.6 Estimation of uranium removal

Uranyl nitrate [UO_2_ (NO_3_)_2_.6H_2_O] solution (1000 mg/L) was prepared in ultrapure water and filter sterilized. The pH of the aqueous solution was 3.61. DR1-bf^+^ biofilms were grown as described above [section 2.4] for the prescribed time of growth as per the experiment plan. TGY medium along with planktonic cells was aspirated from the petri dish and discarded. Biofilm was gently washed twice with sterile PBS. Only TGY medium with Ca^2+^ and kanamycin (excluding bacterial cells) was also included as blank. Washed biofilms were treated with 1000 mg/L of Uranyl nitrate solution. Sample was collected at different time intervals up to 2 hours for the estimation of uranium as per above described method.

### 2.7 Scanning Electron Microscopy (SEM)

DR1_bf^+^ biofilm was grown as explained in the section 2.1 on the glass slides without agitation. Biofilm was fixed with 2.5% glutaraldehyde (in PBS, pH 7.2) at 4°C overnight. The biofilm was scrapped out, washed with PBS thrice and dehydrated with a series of increasing ethanol solutions (10, 20, 30, 40, 50, 60, 70, 80, 90 and 100 % w/v, with 10 minutes incubations at each step). The slides were coated with gold and were analysed using scanning electron microscopy (HELIOS SEG-SEM). Analysis of SEM images revealed the presence of dense biomass embedded in EPS matrix.

### 2.8 Fourier transform infrared spectroscopy (FTIR)

The 4-day old DR1bf^+^ biofilm was grown as explained in (section 2.4) and scrapped out (before and after treating with uranium) in to an Eppendorf. The biomass was lyophilised and taken for FTIR analysis. The infrared spectra of lyophilised bacterial biomass samples (before and after uranium treatment) were recorded within a range 400–4,000 cm^-1^ in an ABB MB3000 FTIR Spectrometer. The samples were prepared as KBr pellets.

## 3. Results

### 3.1 Standardisation of uranium estimation protocol

Figure 1 shows that the arsenazo method gave a linear correlation up to 20 mg/L concentration of uranium with 0.01% arsenazo II dye solution using potassium hydrogen phthalate buffer. The concentrations estimated by this method were cross checked for its accuracy by using ICP-AES analysis, the values corroborated with the modified arsenazo III method. By changing the concentration of arsenazo III dye the concentration estimated can go up to 1000 mg/L. Thus, this modified arsenazo method was used for uranium estimation in this study.

**Figure 1:**
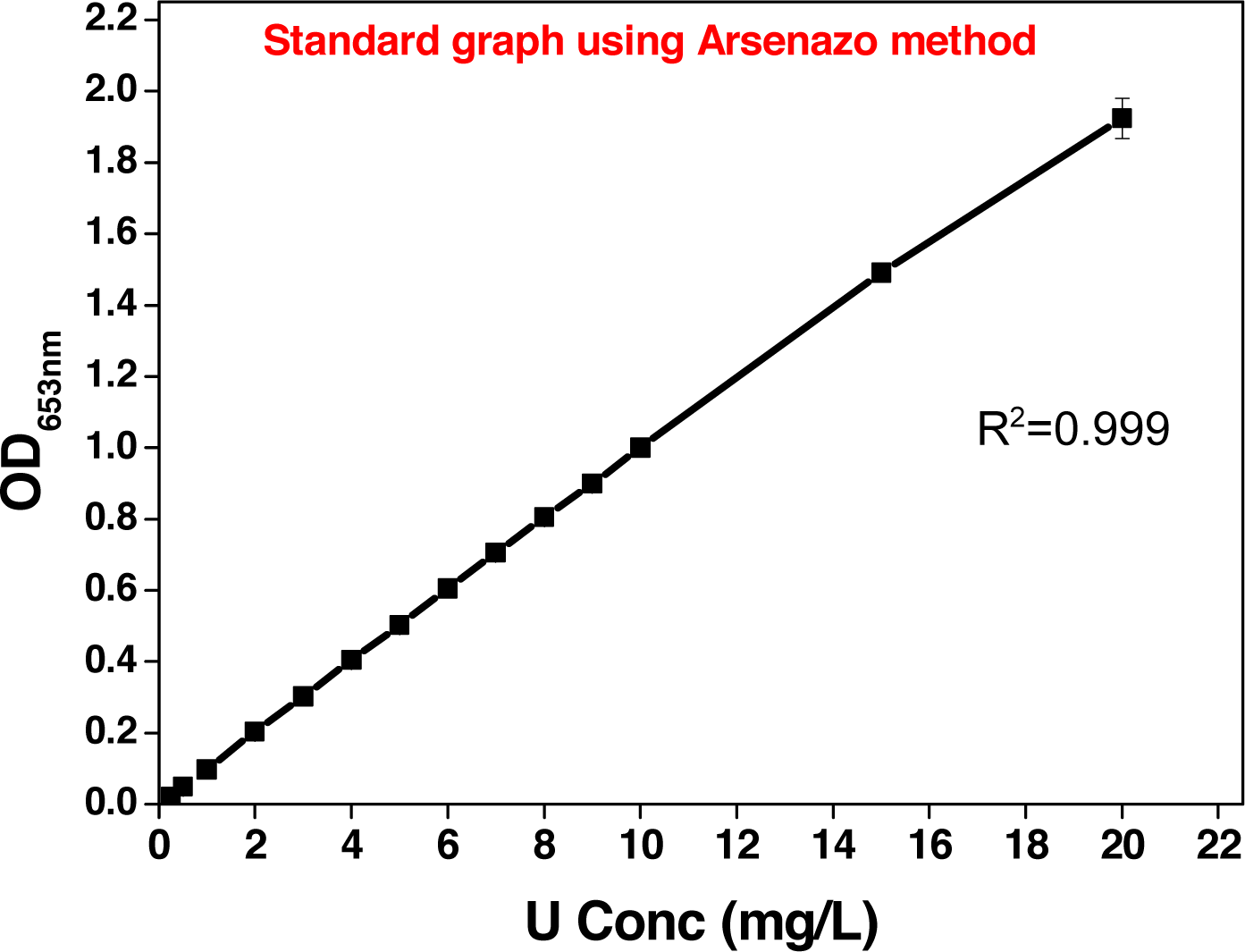
Standard graph showing linear correlation between concentration of uranyl ion and absorbance at 653 nm using modified Arsenazo II dye method.

### 3.2 Biofilm growth by *D. radiodurans* R1 (DR1-bf^+^) strain

The crystal violet assay showed that the recombinant *D. radiodurans* (DR1-bf^+^) has a very good ability to form biofilm in contrast to its wild type counterpart (Figure 2a). Fluorescence microscopic study also confirmed the biofilm formation (Figure 2b). The maximum biofilm formation was observed at 4 days and calcium in the growth medium enhanced the biofilm production in DR1-bf^+^ proportionally. This shows the effect of Ca^2+^ on EPS production and relative increase in thickness of biofilm (Shukla and Rao, 2015). 20 mM concentration of Ca^2+^ was shown to have optimum effect on the extracellular matrix production of DR1-bf^+^ biofilm compared to other concentrations of calcium ion (Shukla and Rao, 2017a). Therefore this concentration of Ca^2+^ was used for rest of the experiments.

**Figure 2:**
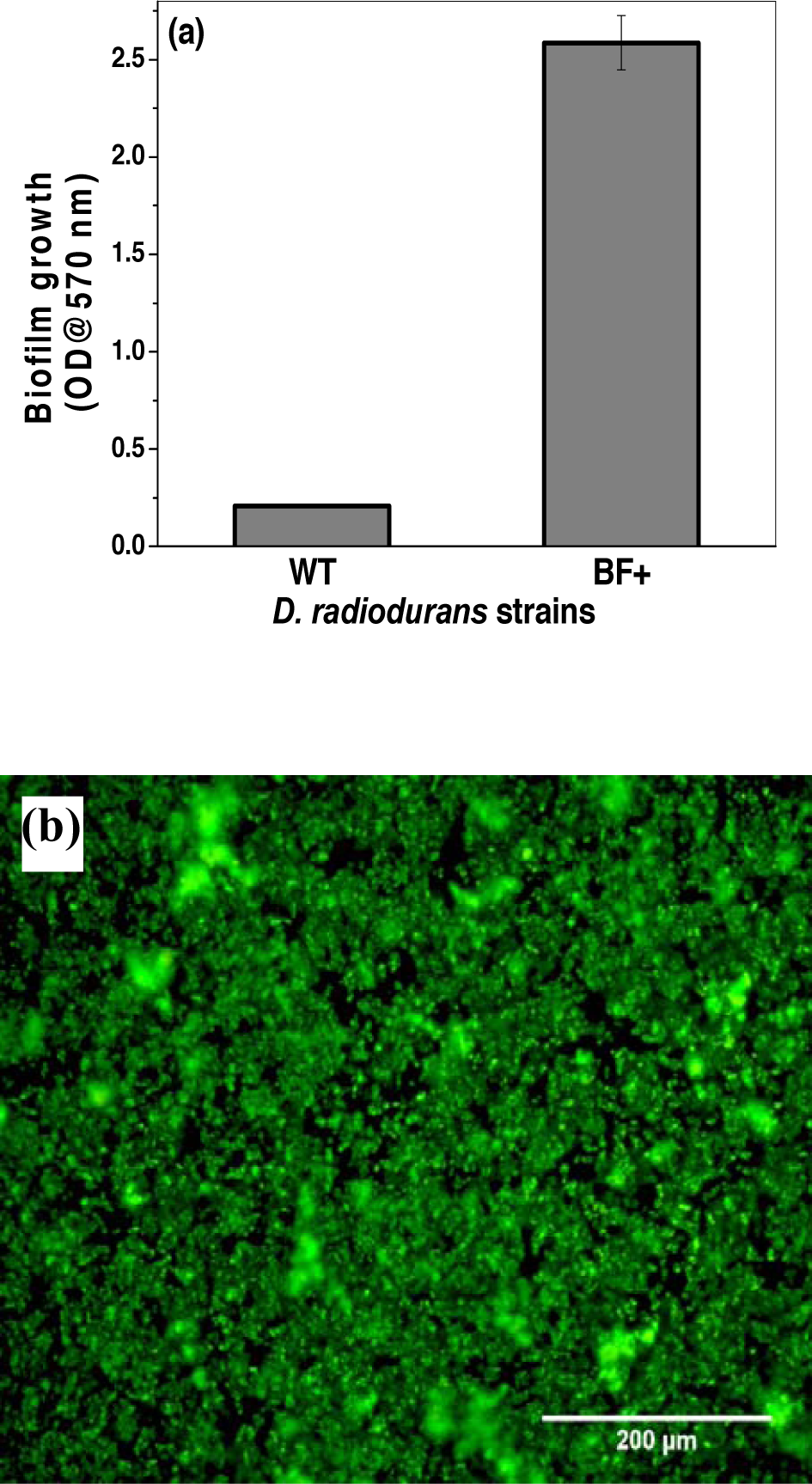
Biofilm growth by *D. radiodurans* (DR1-bf^+^) strain. (a) CV assay conforming the biofilm growth in the recombinant biofilm strain (DR1-bf^+^) compared to the wild type. (b) Florescence microscope image showing the attachment of DR1-bf^+^ cells, forming a biofilm. DR1-bf^+^ showing the green fluorescence due to the presence of constitutively expressing GFP.

### 3.3 Bioremediation of uranium using DR1-bf^+^ biofilm and planktonic cells of DR1-WT and DR1-bf^+^

The biofilm was grown for 4 days and treated with aqueous uranium solution as described in the sections 2.4 and 2.6. A control biofilm grown in the absence of Ca^2+^ was also evaluated for comparison. Correspondingly, the planktonic cells, both of *D. radiodurans* R1 wild type (DR1-WT) and *D. radiodurans* (DR1-bf^+^) strains were grown in test tubes, maintaining the identical growth conditions but in the absence of Ca^2+^. The assay was performed after 4 days of biofilm growth. Required amount of sample was collected at different time intervals and the reduction in uranium concentration was estimated. Results showed that the uranium remediation by DR1-bf^+^ biofilm in the presence of Ca^2+^ was very quick in the first 5 min post treatment, which removed nearly 25% of initial concentration of uranium (1000 mg/L). Beyond which the removal rate slowly decreased and reached a maximum of 75 ± 2% removal within 30 min of the treatment. With increase in time, further removal of uranium was not observed, which might be due to saturation of its adsorption capacity of the biofilm.

As shown in the Figure 3A (by Arsenazo dye method) and Figure 3B (by ICP-AES) removal of uranyl ion by DR1-bf^+^ biofilm when grown in the presence of Ca^2+^, was significantly high as compared to DR1 planktonic cells, DR1-bf^+^ planktonic cell or DR1-bf^+^ control biofilm (grown without Ca^2+^). Results showed that the removal efficiency of DR1-bf+ (+ Ca^2+^) biofilm was four times more as compare to DR1-bf+ planktonic cells. A visible yellow precipitate of uranyl nitrate by the biofilm was observed after the remediation process (Figure 3a) that indicates the conversion of aqueous form of U(VI) to an insoluble precipitate U(IV), making the removal of uranium more competent than other methods. Although initial experiments showed that the mechanism of uranium bioremediation appears to be an adsorption phenomenon, the possibility of enzyme mediated process may not be ruled out, at this stage of study.

**Figure 3:**
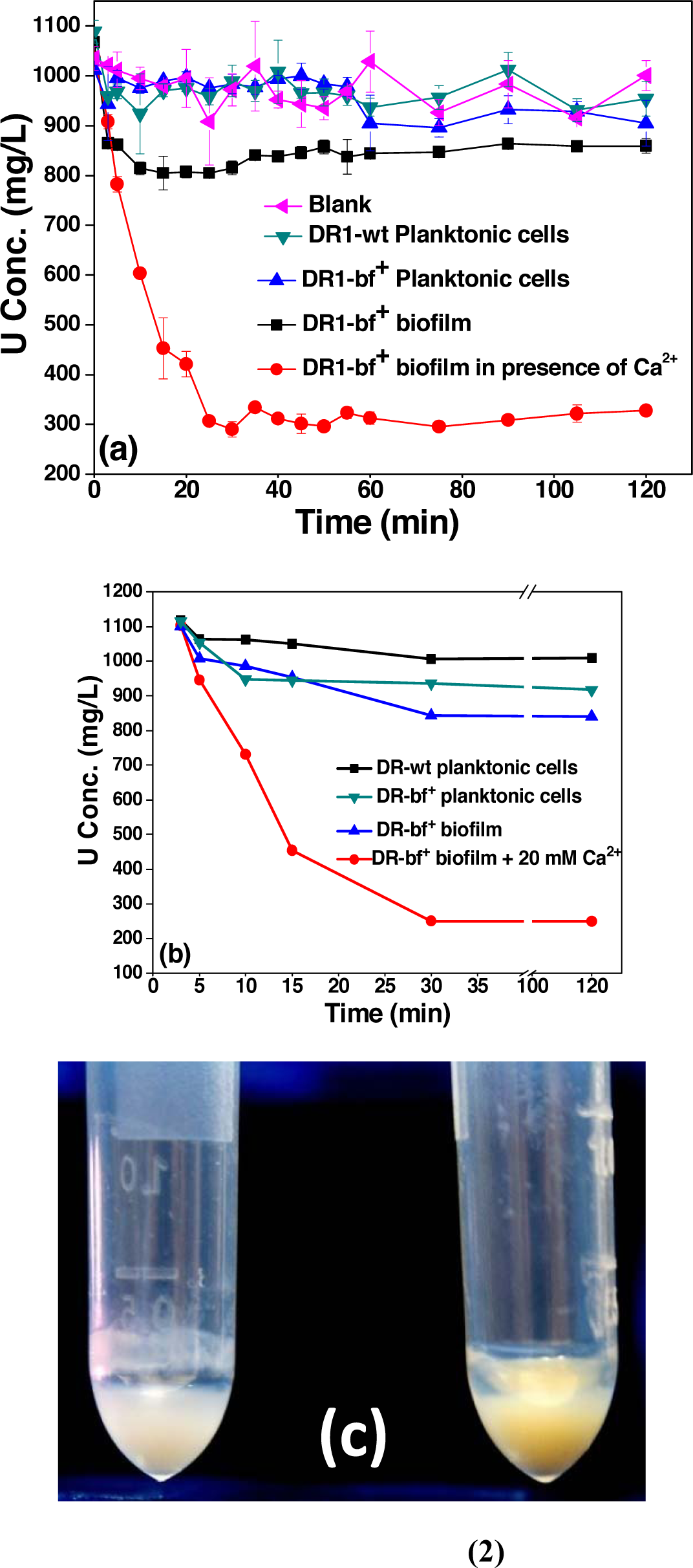
Comparison of uranium removal by *D. radiodurans* biofilm (grown with or without Ca^2+^) and planktonic cells of *D.radiodurans*-wild type and DR1-bf^+^. Uranium concentrations were estimated by (A) Arsenazo dye method and (B) ICP – AES analysis. (C) Visible precipitation of uranium (yellow colour) was observed in the biofilm. Control DR1-bf^+^ biofilm biomass (2) DR1-bf^+^ biofilm biomass treated with uranium aqueous solution (1000 mg/L) for 5 min.

### 3.4 The effect of biofilm age on uranium bioremediation

To study the effect of biofilm’s age on the efficiency of uranium removal, a day-wise U removal assay was performed. Results showed that there was a continuous increase in uranium removal efficiency as the DR1-bf^+^ (+Ca^2+^) biofilm grew older (see Figure 4). The removal rate was found to be higher after 4 days of biofilm growth. Without much increase in biofilm biomass, this suggests that there is a continuous change in the biofilm matrix composition over a period of time (Table 1).

**Table 1:** Table showing the amount of biomass produced with the increse in the age of the bioiflm. These biofilm were grown as described in section 2.4. The biofilm was scrapped and dried to get the weight.

| Biofilm's age | Biofilm biomass [mg] |
| --- | --- |
| Day 1 | 29.24 |
| Day 2 | 29.97 |
| Day 3 | 29.43 |
| Day 4 | 27.38 |
| Day 5 | 29.38 |
| Day 6 | 28.03 |
| Day 7 | 32.43 |

**Figure 4:**
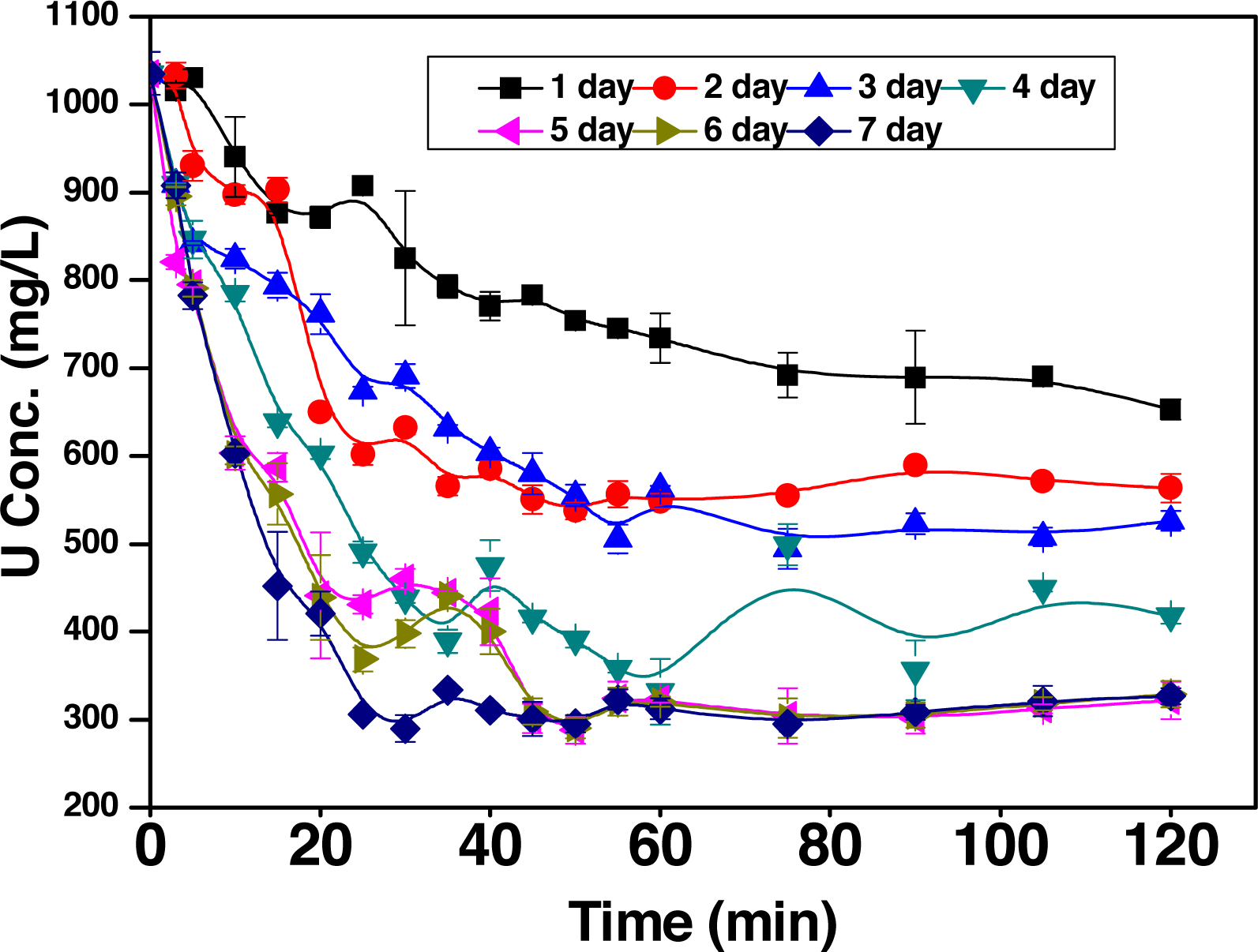
Uranium removal by different day old DR1-bf^+^ biofilm grown in the presence of 20 mM Ca^2+^

**Figure 5:**
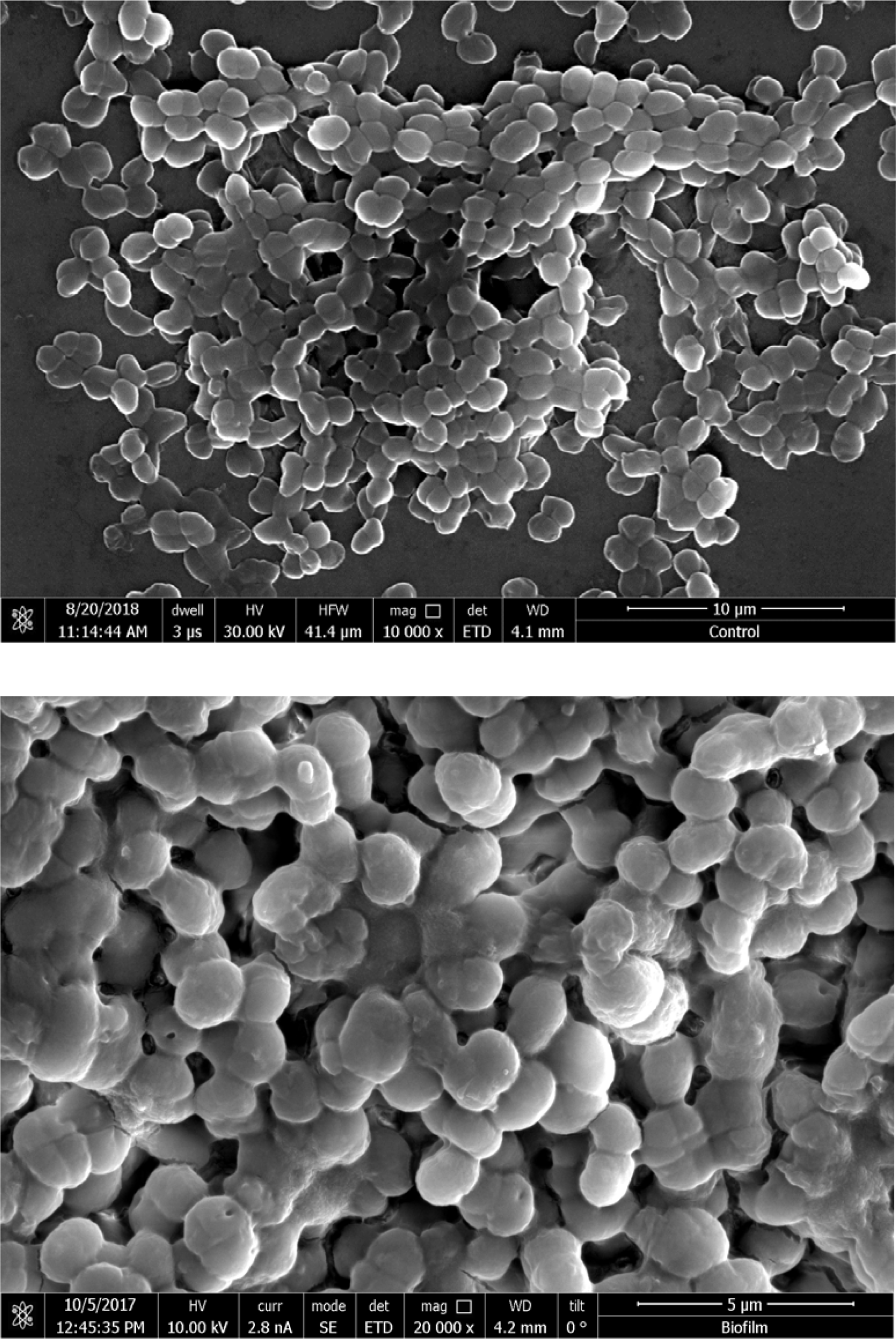
Scanning Electron Microscopic images of *Deinoccoccus radiodurans* (DR1_bf^+^) biofilm.

Our previous studies showed that there was an enhanced exopolysaccharide and eDNA production in the biofilm matrix in the presence of Ca^2+^ (Shukla and Rao, 2014; Shukla and Rao, 2015). In view of this, it is speculated that uranyl ion (UO_2_^2+^) removal is by biosorption process where negatively charged exopolysaccharide and eDNA play a critical role. However, visible precipitation and faster removal of UO_2_^2+^ by older biofilm also suggest that this phenomenon can also be aided by enzymatic reduction.

### 3.5 Adsorption is the major method of uranium sequestration

FTIR spectroscopic study for the control biofilm (uranium-free) and the uranium treated biomass were performed to elucidate the chemical groups involved in uranium binding (Figure 6). The highly complex IR spectra was analysed to assign the important characteristic peaks which illustrates the involvement of main functional groups present in the bacterial biomass. The amino group (N–H stretching) peak lies in the spectrum region occupied by a broad and strong band (3,600–3,000 cm^-1^). This may be due to hydroxyl groups that are hydrogen bonded to various degree (Pagnanelli et al., 2000). In the control spectrum, absorption peaks at 2,900–3,000 cm^-1^ are ascribed to the asymmetric stretching of C–H bond of the –CH_2_ groups combined with that of the CH_3_ groups. In the control spectrum, the γC=O of amide I and δBNH/γC=O combination of the amide II bond were present at 1,647 and 1,546 cm^-1^, respectively, indicating the presence of carboxyl groups (Kazy et al., 2006). In the uranium loaded samples, these peaks were shifted from 1,647 cm^-1^ to 1,650 cm^-1^ and 1,546 cm^-1^ to 1,540 cm^-1^ region, indicating an interaction between uranium and carboxyl group. Especially, 1540 cm^-1^ peak indicates the scissoring of CH_2_ group. Sharp spectra from 1456-1481 cm^-1^ ascribe to carboxyl group. A decrease in the intensity and a gradual shift of the peak at 1214 cm^-1^, in the control sample to a lower energy in uranium (1204 cm^-1^) loaded samples clearly shows the weakening of the P = O as a result of uranium binding to the phosphates. A sharp peak at 1387 cm^-1^ is indicative of symmetrical bending of CH_3_ group (Stuart, 2005). In uranium loaded sample, distinct peak at 927 cm^-1^ and changes in peak positions and intensity around 1000 cm^-1^ region can be assigned to asymmetric stretching vibrations of γ3 UO_2_^2+^ and stretching vibrations of weekly bounded oxygen ligands with uranium (γ U-O ligand) (Kazy et al., 2009). The overall spectral analysis strongly supports the major role of carboxyl, amide and phosphate groups in uranium binding by the bacterial biomass.

**Figure 6:**
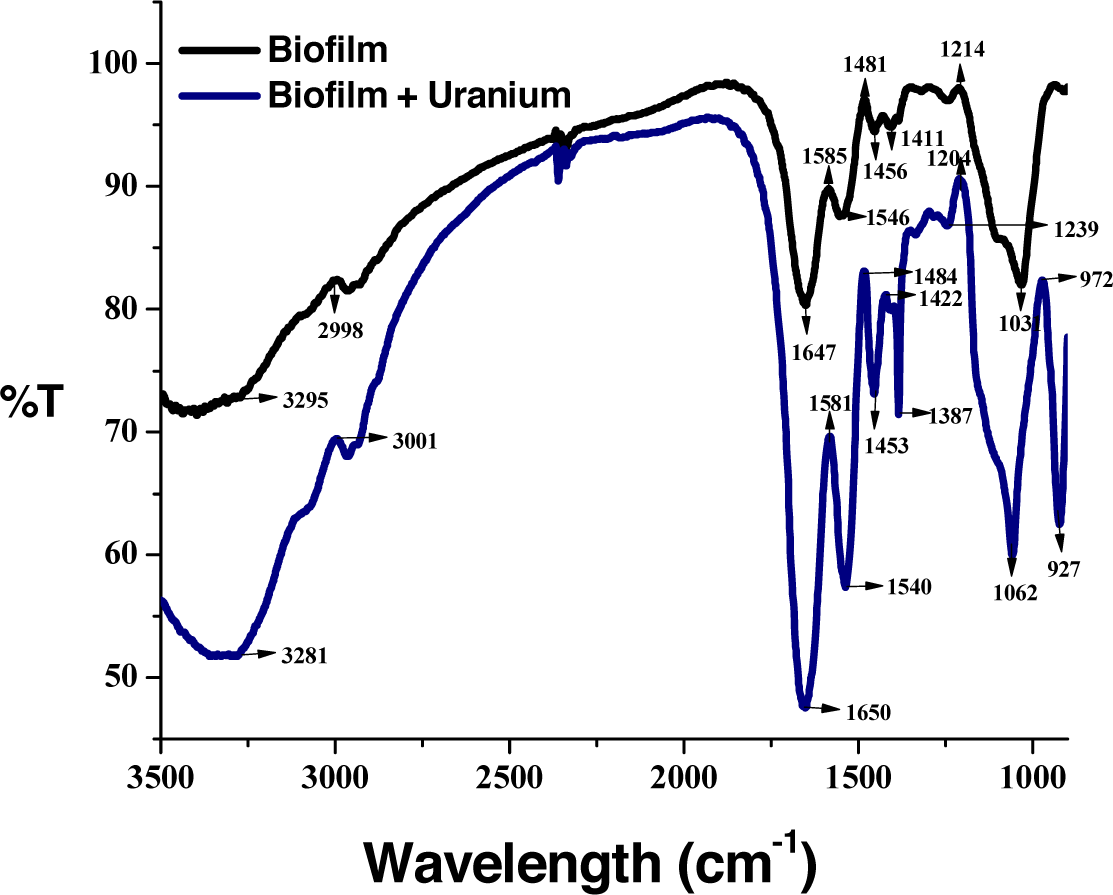
Fourier transformed IR spectra of *D. radiodurans* (DR1_bf^+^) biofilm biomass: before (a), and after uranium treatment (b).

## 4. Discussion

In the present study the use of a recombinant *D. radiodurans* biofilm for the bioremediation of uranium from aqueous solution showed remarkable results. Although planktonic forms of these species have been used for some treatment process, their drawbacks restrict their use for a wider application. For example, earlier studies involving U remediation using *D. radiodurans,* showed good removal efficiency up to 95% when treated with uranium concentration ranging from 5-100 µM, with an incubation period of 21 days, which greatly affected the efficiency of the system (Fredrickson et al., 2000). Though in few studies involving few bacterial cultures such as *Desulfosporosinus orientis* DSM 765, *Desulfosporosinus spp.* P3 (Suzuki et al., 2004), *Clostridium sp.* (Francis et al., 1994) and *Clostridium sphenoides* ATCC 19403 (Francis et al., 2004) have showed a significant uranium removal under anaerobic environment. However, maintaining the anaerobic condition is another challenging task and a major bottle neck while applying at industrial scale. Marsili et al., (2005) reported the use of sulphate reducing bacterial biofilm for uranium removal with preformed biofilm grown for 3 weeks in a bioreactor. The efficiency of removal was estimated to be 30.4% in the reactor supplied with 3 mg/L of U and 73.9% in the reactor supplied with 30 mg/L of U. The reactor performance was not up to the expectation, which was hypothetically ascribed to the differences in biofilm sloughing events in the reactor. Although these reports were evident of uranium removal process by bacteria, none of the systems has been proved to be efficient enough for the large-scale applications. Application of such bacteria in U bioremediation (low active waste) will be further difficult as they do not have the capability to survive in the presence of radioactivity.

*Deinococcus radiodurans* is known for its extreme radiation tolerance (Krisko and Radman, 2013). *D. radiodurans* does not have innate ability to form biofilm. Although *Deinococcus geothermalis* (Kolari et al., 2002) has shown the ability to form biofilm, no study has indicated its potential application in bioremediation. This study involves the use of a genetically modified *D. radiodurans biofilm* for uranium bioremediation purpose. The genetically modified strain of *D. radiodurans* (DR1-bf^+^), when grown in the presence of 20 mM Ca^2+^ was found to produce very good biofilm thus DR1-bf^+^ biofilm, grown in the presence Ca^2+^ was tested for the uranium bioremediation in this study. Ca^2+^ plays an important role in the adhesion of bacterial cells to the surfaces. They are involved both in specific (actions which cannot be replaced by other cations) and non-specific interactions (between cells and substratum) in biofilms (Shukla and Rao, 2013; Mangwani et al., 2014). Also, the interaction of protein and polysaccharide at the cell surface is influenced by Ca^2+^ (Geesey et al., 2000). In the biofilm, extracellular Ca^2+^ ions involve in the cross-linking of extracellular polymeric substances, which gives strength to the biofilm matrix and regulates production of many extracellular enzymes. It also plays an important role in maintaining the integrity of the biofilm (Shukla and Rao, 2014). In a way, Ca^2+^ acts as an important ionic cross-bridging molecule for negatively charged bacterial polysaccharides. It was also shown in our previous study that production of extracellular matrix was enhanced in *D. radiodurans* biofilm when biofilm was grown in the presence of Ca^2+^ in the growth medium as compared to that of biofilm grown in the absence of Ca^2+^ (Shukla and Rao, 2017a). Among the various concentrations of Ca^2+^ used in the study, 20 mM showed a significant improvement in the biofilm production, as compared to other concentrations (Shukla and Rao, 2017a). It is possible that the Ca^2+^ can influence the *D. radiodurans* biofilm growth in multiple ways but as of now, the exact mechanism is not clear. The involvement of biosorption by negatively charged exopolysaccharide and eDNA in the biofilm matrix of *D. radiodurans* has been speculated to be the reason for this remediation process also the visible precipitate formation and detection of acid phosphatase activity cannot rule out the possibility of enzyme-mediated reduction of U(VI) to U(IV) (Francis et al.).

Our results showed that 4 day old DR1-bf^+^ biofilm had significant capability to remove UO_2_^+2^ ions that too at a very high rate which was ∼75% removal within 30 mins (Figure 3A). This observation implicates the potential of *D. radiodurans* for the development of biofilm-based bioremediation process for UO_2_^2+^ removal from radioactive aqueous waste solutions. The biofilm mode of *D. radiodurans* has shown tolerance to high concentrations of uranium, up to 1000 mg/L. The DR1-bf^+^ biofilm could withstand very high concentrations of uranyl nitrate solution i.e. 1000 mg/L not only due to the fact that *D. radiodurans* has significant heavy metal tolerance (Brim et al., 2000) but also of the fact that biofilm mode of life reduces the toxicity of heavy metals to microbes as compared to that of planktonic cells. This DR1-bf^+^ biofilm-mediated uranium removal method showed significant higher efficiency in terms of both performance as well as time (75% uranium removal within 30 min post-treatment) as compared to its planktonic counterparts as well as *D. radiodurans* wild type strain. No requirement of special conditions such as maintaining an anaerobic condition in this removal process and tolerance to high concentrations of uranium make *D. radiodurans* biofilm-mediated U removal process promising. Further investigations into the form of uranium which is precipitated and the mechanism involved in the microbial reduction of this U has to be evaluated. These studies may provide insights into strategies for promoting the microbial reduction in uranium-contaminated sites.

## 5. Conclusion

*Deinococcus radiodurans* (DR1-bf^+^) biofilm was tested for its potential to remove uranium from aqueous solution. The study showed that when DR1-bf^+^ is grown in the presence of 20 mM Ca^2+^, it acquired an extraordinary capacity of UO_2_^+2^ removal as compared to that of its planktonic counterparts and control biofilm. In this study, it was shown that 4 day old DR1-bf^+^ biofilm when grown in the presence of 20 mM Ca^2+^, significantly removed UO_2_^+2^ ions (∼75% removal) within few minutes. There was an increase in the uranium removal capacity with increase in the age of the biofilm. Interestingly, there was not much variation in the biofilm biomass with the age. Without significant change in the biofilm biomass, age-dependent increase in U bioremediation was observed. FTIR analysis strongly supports the involvement of carboxyl, amide and phosphate groups in uranium binding by the bacterial biomass. This observation suggest that uranyl ion (UO_2_^2+^) removal by DR1-bf^+^ is a biosorption process, where negatively charged exopolysaccharides and eDNA play critical roles. This study implicates the potential of *D. radiodurans* biofilm-based bioremediation process for UO ^2+^ precipitation.

